# Mechanism and resistance for antimycobacterial activity of a fluoroquinophenoxazine compound

**DOI:** 10.1101/464990

**Authors:** Pamela K. Garcia, Thirunavukkarasu Annamalai, Wenjie Wang, Raven Bell, Duc Le, Paula Martin Pancorbo, Sabah Sikandar, Ahmed Seddek, Xufen Yu, Dianqing Sun, Anne-Catrin Uhlemann, Purushottam B. Tiwari, Fenfei Leng, Yuk-Ching Tse-Dinh

**Affiliations:** Department of Chemistry & Biochemistry, Florida International University, Miami, Florida, United States of America; Biomolecular Sciences Institute, Florida International University, Miami, Florida, United States of America; Department of Pharmaceutical Sciences, The Daniel K. Inouye College of Pharmacy, University of Hawaii at Hilo, 34 Rainbow Drive, Hilo, Hawaii, United States of America; Division of Infectious Diseases, Columbia University Medical Center, New York, New York, United States of America; Department of Oncology, School of Medicine, Lombardi Comprehensive Cancer Center Georgetown University Medical Center, Washington DC, United States of America

## Abstract

We have previously reported the inhibition of bacterial topoisomerase I activity by a fluoroquinophenoxazine compound (FP-11g) with a 6-bipiperidinyl lipophilic side chain that exhibited promising antituberculosis activity (MIC = 2.5 μM against *Mycobacterium tuberculosis*, SI = 9.8). Here, we found that the compound is bactericidal towards *Mycobacterium smegmatis*, resulting in greater than 5 Log_10_ reduction in colony-forming units [cfu]/mL following a 10 h incubation at 1.25 μM (4X MIC) concentration. Growth inhibition (MIC = 50 μM) and reduction in cfu could also be observed against a clinical isolate of *Mycobacterium abscessus.* Stepwise isolation of resistant mutants of *M. smegmatis* was conducted to explore the mechanism of resistance. Mutations in the resistant isolates were identified by direct comparison of whole-genome sequencing data from mutant and wild-type isolates. These include mutations in genes likely to affect the entry and retention of the compound. FP-11g inhibits *Mtb* topoisomerase I and *Mtb* gyrase with IC_50_ of 0.24 and 31.5 μM, respectively. Biophysical analysis showed that FP-11g binds DNA as an intercalator but the IC_50_ for inhibition of *Mtb* topoisomerase I activity is >10 fold lower than the compound concentrations required for producing negatively supercoiled DNA during ligation of nicked circular DNA. Thus, the DNA-binding property of FP-11g may contribute to its antimycobacterial mechanism, but that alone cannot account for the observed inhibition of Mtb topoisomerase I.

## Introduction

Tuberculosis (TB) is a devastating disease caused by *Mycobacterium tuberculosis* infection. The World Health Organization (WHO) reported that TB is the leading cause of death worldwide from a single infectious agent. The 2018 Global Tuberculosis Report estimated that TB caused ~1.6 million deaths in 2017, with up to 10 million newly diagnosed TB cases annually [1]. Significantly, drug-resistant TB poses a great threat to public health. In 2017, there were 558,000 new cases of drug resistant TB, including multidrug resistant TB (MDR-TB). resistant to rifampicin and isoniazid (two first-line anti-TB drugs) and rifampicin-resistant TB (RR-TB) [1]. These TB patients are more likely to have poor treatment outcomes. Therefore, there is an urgent need for potent new TB drugs with novel modes of action [2]. Resistance to current antibiotics also makes it difficult to treat infections caused by nontuberculous mycobacteria (NTM). It is often difficult to distinguish between *M. tuberculosis* and NTM as the cause of lung infections, so there is a medical need for new antibiotics that have broad antimycobacterial activity.

DNA topoisomerases are essential enzymes for maintaining optimal DNA topology to facilitate vital functions including replication, transcription, recombination and DNA repair [3]. Topoisomerases belong to type I or type II families corresponding to cleavage of a single or double strand of DNA being utilized for catalysis [4] with subfamilies based on sequence homology and mechanism [5]. Every bacterium has at least one type IA topoisomerase for overcoming topological barriers that require cutting and rejoining of a single strand of DNA [3, 6]. Topoisomerase I catalytic activity is essential for the viability of *Mycobacterium smegmatis* (*Msm*) and *M. tuberculosis* (*Mtb*) [7, 8], because these organisms only have one type IA topoisomerase. *Mtb* topoisomerase I (MtbTopI) is a validated anti-TB target; [7, 9, 10] for identification of inhibitors that can potentially be developed into leads for new TB therapy.

We have previously shown [11] that a fluoroquinophenoxazine compound (FP-11g, shown in Scheme 1) with a 6-bipiperidinyl lipophilic side chain inhibited the catalytic activity of *Escherichia coli* topoisomerase I (IC_50_ = 0.48 μM), and showed promising antituberculosis activity (MIC = 2.5 μM against *Mtb* H37Rv, SI = 9.8 for Vero cell cytotoxicity). In this study, we followed up on the antimycobacterial activity of this compound to further explore its therapeutic potential. The interactions of this compound with MtbTopI and DNA were characterized in biophysical assays. Mechanism of resistance to its antimycobacterial activity was studied with *M. smegmatis.* This commonly used model mycobacterial organism is non-pathogenic and fast growing, but still shares many features with pathogenic mycobacteria.

## Materials and Methods

### Synthesis of FP-11g

FP-11g (Scheme 1) was synthesized using our previously reported procedure [11]. The structure was confirmed by ^1^H NMR, and high-resolution mass spectrometry (HRMS). Purity was determined to be >99% by reverse phase C18-HPLC.

### MIC (minimum inhibitory concentration) against *M. smegmatis* and *M. abscessus*

*M. smegmatis* mc2 155 (WT from ATCC) and FP-11g resistant mutants derived from WT were cultured in 5 ml Middlebrook 7H9 broth containing 0.2% glycerol, 0.05% Tween 80 with or without a supplement of 10% albumin, dextrose, sodium chloride (ADN) at 37°C with shaking. Stationary phase bacteria cultures were adjusted to an optical density (OD_600_) of 0.1 and subsequently diluted 1:10 using growth media. Fifty microliters (~10^5^ cfu) of the diluted culture were transferred to the individual wells of a clear round-bottom 96-well plate containing 50 μl of serially diluted compounds. The 96-well plate was then incubated at 37°C with shaking. After 48 hours of incubation, resazurin (final concentration 0.002%) was added to the individual wells and the fluorescence reading at 560/590 nm was taken with a BioTek Synergy plate reader after approximately 5h of incubation at 37°C.

A clinical isolate of *M. abscessus* bacterium (isolated at the Columbia University Medical Center) was cultured in Middlebrook 7H9 ADN broth till it reached an optical density (OD_600_) of 1.0. The culture was stored at −80°C as 1 ml aliquots containing 15% glycerol. These frozen aliquots were then used as the inoculum for *M. abscessus* MIC assays. Prior to conducting each MIC assay, an aliquot of frozen *M. abscessus* was thawed and diluted 1:100 in Middlebrook 7H9 broth. After dilution, the bacterial cells (~10^5^ cfu) were added to the wells of a 96-well plate containing the serially diluted compounds as described for *M. smegmatis* cells and incubated at 37°C with no shaking for 48 h. Subsequently, resazurin (final concentration 0.005%) was added to each well and fluorescence reading at 560/590 nm (excitation/emission) was taken after 24 h of incubation at 37°C. MIC was determined as compound concentration that showed at least 90% reduction in fluorescence when compared to the control (no inhibitor) wells. Ciprofloxacin and Clarithromycin were included in the assay as positive controls. On the day of each MIC assay, thawed inoculum of *M. abscessus* was also serially diluted and spread plated on LB agar plates to confirm for the inoculum load of ~10^5^ cfu/well. MIC determination for each bacteria was repeated at least three times.

### Survival assays

The bactericidal effect of FP-11g compound was evaluated in 96-well plates using a protocol similar to the MIC assay. In brief, *M. smegmatis* was incubated at 37°C with shaking in the presence of 1X, 2X, 4X and 8X MIC of FP-11g for 6, 10, 24 and 48 hours; *M. abscessus* was incubated at 37°C without shaking in the presence of 1X and 2X MIC of FP-11g for 24, 48 and 72 hours. At each time point 20 μl from the treatment wells were serially diluted (10 fold), spread on LB agar plates and incubated at 37°C for 4 or 7 days for counting the viable colonies of *M. smegmatis* and *M. abscessus* respectively. Ten microliters from the treatment wells were also enriched in 5-ml of Middlebrook 7H9 broth if there are no viable colonies from a particular treatment-time combination on the LB agar plates. The survival percentage was calculated by dividing the number of viable colonies at each time point by the initial viable count prior to the treatment (time 0). Survival assays were repeated three times.

### Isolation of resistant mutants

*M. smegmatis* mc2 155 was exposed stepwise to increasing concentrations of FP-11g to isolate mutant strains with different levels of resistance for FP-11g by a slightly modified protocol as described by Fujimoto-Nakamura et al [12]. Overnight culture of *M. smegmatis* was adjusted to an optical density (OD_600_) of 0.1 and a volume of 100 μl (2×10^6^ cfu) was spread on 7H9 ADN agar containing 2.5 μM (8X MIC) FP-11g for isolation of resistant mutants. Colonies appearing on these plates were scored as resistant mutants only if they grew again on 7H9 ADN broth and agar plates containing 8X MIC of FP-11g. Mutation frequency was calculated based on these resistant colonies. Only one resistant mutant was isolated at 8X MIC of FP-11g. This resistant mutant was next exposed to 16X MIC of FP-11g in 7H9 ADN broth to further isolate resistant colonies PGM1, PGM2 shown in Table 1. As our initial resistant mutation isolation at 8X MIC of FP-11g resulted in only one mutant, a second round of stepwise mutation was initiated with 4X MIC to higher concentrations of FP-11g (PGM3 – PGM6 shown in Table 1).

**Table 1.** FP-11g resistant mutants, resistance levels and cross-resistance to Moxifloxacin

| *M. smegmatis*<br>Strain | FP-11g<br>concentration for<br>mutant isolation<br>X MIC (0.31 μM) | FP-11g<br>MIC (μM) | Fold-increase in<br>FP-11g MIC | Moxifloxacin<br>MIC (μg/ml) | Fold-increase in<br>Moxifloxacin<br>MIC |
| --- | --- | --- | --- | --- | --- |
| WT | N/A | 0.31 | N/A | 0.08-0.16 | N/A |
| PGM1 | 8X → 16X | 2.5 | 8X | 0.16 | 2X |
| PGM2 | 8X → 16X | 5 | 16X | 0.32 | 4X |
| PGM3 | 4X → 10X | 5 | 16X | 0.32 | 4X |
| PGM4 | 4X → 13X | 10 | 32X | 0.32 | 4X |
| PGM5 | 4X → 16X | 5 | 16X | 0.32 | 4X |
| PGM6 | 4X → 16X | 5 | 16X | 0.16 | 2X |

### Whole genome sequencing

*M. smegmatis* WT and mutant strains were cultured in 5 ml of Middlebrook 7H9 ADN broth and 2 ml was spun down for genomic DNA extraction. DNA extraction was performed using the BACTOZOL^TM^ Bacterial DNA isolation kit (Molecular Research Center) according to manufacturer’s instructions. DNA quality was evaluated through the UV absorbance 260/280 ratio (>1.8) and DNA quantity was measured using a fluorescence-based Qubit^®^ dsDNA BR (Broad-Range) Assay Kit (Thermo Fisher). DNA concentration was adjusted to 0.2 ng/μl with 10 mM Tris-HCl pH 8.0. After concentration adjustment, all the genomic DNA samples were quantified once again to confirm the concentration. Library was prepared with the Nextera XT DNA Library Prep Kit, 96 Indexes (FC-131-1096). After clean-up of the libraries, random samples were selected for the Bio-analyzer analysis to verify the average size of the fragments. Cleaned up libraries were normalized manually and pooled together. The pooled libraries were then denatured with NaOH (final concentration 0.1 N) and the PhiX Control v3 library added. This mix was loaded in the pre-thawed reagent cartridge associated with the MiSeq Reagent kit v3 and sequenced using Illumina MiSeq^®^ Next Generation Sequencer. FASTQ files generated from the sequencing run was further analyzed in CLC genomics workbench 10 software (QIAGEN). Sequence of *M. smegmatis* mc2 155 WT strain from our laboratory was used as reference to compare all the mutant sequences and detect variations (Single Nucleotide Variations, Deletions and Insertions).

### Enzyme assays

*MtbTopI relaxation inhibition assay* - MtbTopI was expressed in *E. coli* T7 Express crystal strain (New England Biolabs) and purified as previously described [13,14]. The relaxation activity of MtbTopI was assayed in a buffer containing 10 mM Tris-HCl, pH 8.0, 50 mM NaCl, 0.1 mg/ml gelatin, and 0.5 mM MgCl_2_. Serial dilutions of FP-11g dissolved in DMSO was mixed with 10 μl of the reaction buffer containing 12.5 ng of enzyme before the addition of 9 μl of reaction buffer containing 160 ng of supercoiled pBAD/Thio plasmid DNA purified by cesium chloride gradient as the substrate. The reactions were terminated following an incubation of the mixtures at 37°C for 30 min by the addition of 4 μl of stop solution (50% glycerol, 50 mM EDTA, and 0.5% (vol/vol) bromophenol blue) and the mixtures were analyzed by agarose gel electrophoresis with TAE buffer (40 mM Tris acetate and 1 mM EDTA, pH 8.2). The gels were stained in ethidium bromide and photographed under UV light.

*DNA gyrase supercoiling inhibition assay* – *Mtb* DNA gyrase was purchased from Inspiralis. The supercoiling assays were carried out by mixing serial dilutions of FP-11g in 0.5 μl DMSO with 1 U of the enzyme in reaction buffer of 50 mM HEPES pH 7.9, 100 mM potassium glutamate, 6 mM magnesium acetate, 4 mM DTT, 1 mM ATP, 2 mM spermidine, 0.05 mg/ml BSA, followed by the addition of 300 ng of relaxed covalently closed circular DNA, for a final reaction mixture volume of 20 μl. The samples were incubated at 37°C for 30 min before the reactions were terminated and analyzed by agarose gel electrophoresis.

### Surface Plasmon Resonance

Biacore T200 SPR instrument was used to conduct all experiments. MtbTopI was used as ligand to immobilize on a CM5 sensor surface in 10 mM sodium acetate buffer at pH 5.5, using Standard amine coupling chemistry, to a level of ~13400 RU. PBS-P (20 mM phosphate buffer, pH 7.4, 2.7 mM KCl, 137 mM NaCl, 0.05% v/v surfactant P20) was used as the immobilization running buffer. FP-11g was used as an analyte to inject over the ligand immobilized surface in various concentrations in the presence of PBS-P buffer supplemented with 10% DMSO. The flow rate of all analyte solutions was maintained at 50 μl/min. One 20 s pulse of 1M NaCl solution was injected for surface regeneration. The contact and dissociation times used were 60s and 300s, respectively. SPR sensorgrams were both reference and bulk subtracted. All experiments were conducted at 25°C.

### Measurement of visible absorbance and fluorescence emission spectra

Visible absorbance spectra of free and bound FP-11g were recorded in a Cary Bio50 UV-VIS spectrophotometer. Fluorescence emission spectra were measured by using a Cary fluorescence spectrophotometer.

### DNA UV Melting studies

DNA UV melting curves were determined using a Cary 100 UV-Vis spectrophotometer equipped with a thermoelectric temperature-controller. The salmon testes (ST) DNA in the presence of different concentrations of FP-11g in 1×BPE buffer (6 mM Na_2_HPO_4_, 2 mM NaH_2_PO_4_, and 1 mM EDTA, pH 7) was used for UV melting studies. Samples were heated at a rate of 1°C min^−1^, while continuously monitoring the absorbance at 260 nm. Primary data were transferred to the graphic program Origin (MicroCal, Inc., Northampton, MA) for plotting and analysis.

### Dialysis experiments

Dialysis assays were carried out according to the previously published procedure [15]. Briefly, a volume of 0.3 ml of 75 μM (bp) of ST DNA was pipetted into a 0.3 ml disposable dialyzer. The dialysis units were then placed into a beaker with 100 ml of 1×BPE buffer (6 mM Na_2_HPO_4_, 2 mM NaH_2_PO_4_, and 1 mM EDTA, pH 7) containing 1 μM of FP-11g. The dialysis was allowed to equilibrate with continuous stirring for 72 hours at room temperature (24°C). After the dialysis, the free, bound, and total concentrations of FP-11g were determined spectrophotometrically. These values were used to determine the DNA binding constant of FP-11g.

### DNA ligation assays

DNA ligation assays were carried out in 1×DNA T4 DNA ligase buffer (50 mM Tris-HCl (pH 7.5), 10 mM MgCl_2_, 1 mM ATP, and 10 mM DTT) using 1,200 units of T4 DNA ligase (New England Biolabs) in 100 μl of solution containing the nicked plasmid pAB1 [16] in the presence of different concentrations of FP-11g. After incubation at 37°C for 30 min, the ligation reactions were stopped by extraction with 100 μl of phenol. The DNA samples were analyzed by electrophoresis in a 1% agarose gel in TAE. After electrophoresis, the agarose gel was stained with ethidium bromide and photographed using a Kodak imaging system.

## Results

### Growth inhibition of *M. smegmatis* and *M. abscessus*

The MIC of FP-11g for *M. smegmatis* mc2 155 was found to be 0.31 μM in multiple measurements. The MIC value is the same in 7H9 media with or without ADN supplement. There is also no effect on the MIC from shaking during the incubation. For the clinical *M. abscessus* isolate, the MIC was 50 μM, compared to Clarithromycin MIC of 0.7-1.56 μg/ml (1-2 μM) and Ciprofloxacin MIC of 12.5 μg /ml (38 μM), IC_50_ for 50% growth inhibition of the *M. abscessus* strain was estimated to be 3-6 μM.

### Bactericidal activity of FP-11g

FP-11g showed strong bactericidal activity against *M. smegmatis.* When *M. smegmatis* cells (~1×10^5^ cells in each well in 96-well plate) were exposed to 2X MIC of FP-11g, the cell viability was diminished by ~ 4.5 Log after 48 hours of incubation (Fig 1A). For concentrations of 4X MIC and 8X MIC, 24 hours were sufficient to diminish cell viability by greater than 5 Log, with no viable cells remaining at 48 hours.

Treatment of *M. abscessus* with 2X MIC of FP-11g resulted in >3 Log loss of cell viability (Fig 1B). The *M. abscessus* strain grows slower than *M. smegmatis*, so treatment was extended to 72 hours. Because of the higher MIC value for *M. abscessus*, FP-11g concentrations could only be tested at up to 4X MIC in the survival assay.

**Fig 1.**
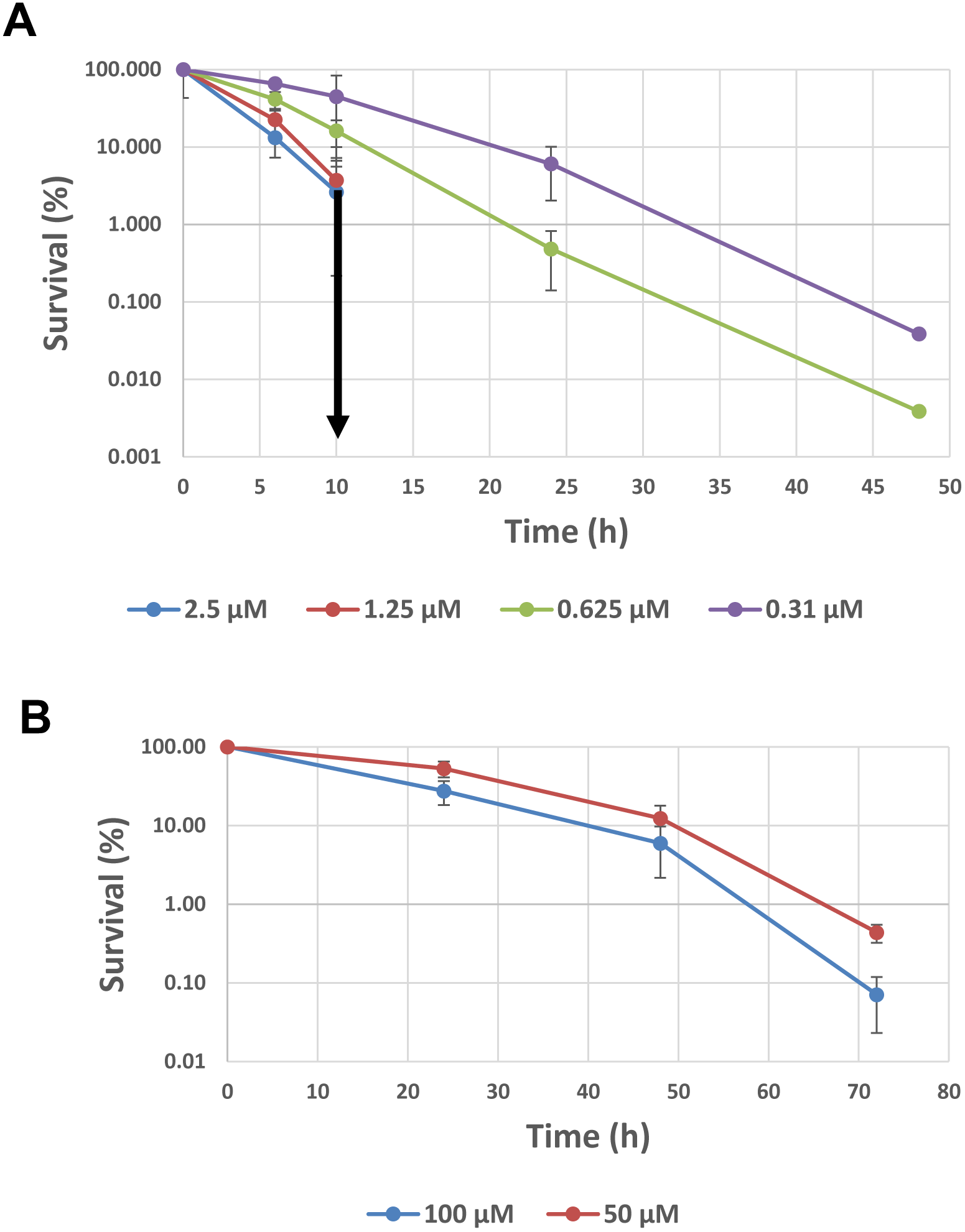
Bactericidal effect of FP-11g on *M. smegmatis* mc2 155 and *M. abscessus*. **(A)** The bactericidal effect of FP-11g on *M. smegmatis* were determined using 1X (0.31 μM), 2X (0.62 μM), 4X (1.25 μM) and 8X (2.5 μM) MIC at different time points. The downward arrow indicates that no viable colonies were detected following treatment with 4X and 8X MIC at time points beyond 10 hours. **(B)** Survival of clinical isolate of *M. abscessus* following treatment with FP-11g at 50 and 100 μM. Percent survival is calculated by the ratio of cfu from treated culture versus cfu from culture prior to addition of FP-11g. The error bars represent the standard deviation of results of experiments repeated three times.

### Isolation and verification of resistant mutants

The mutation frequency for resistance to FP-11g at 8X (MIC) was estimated to be 5×10^−7^ Six *M. smegmatis* strains (PGM1 – PGM6, Table 1) isolated from stepwise increase of FP-11g concentrations were verified to be resistant to the compound by determination of MIC. An increase in the resistance to Moxifloxacin was also observed but the fold-increase in Moxifloxacin was significantly lower than the fold-increase for FP-11g MIC.

### Mutations identified in WGS

The mutations found in each of the resistant strains are listed in Table 2. Mutations in ten different genes associated with deletions, insertions or SNV (Single Nucleotide Variations) were detected in these mutants (Table S1). All of the resistant strains have SNV on *MSMEG*_*0965* gene (*mspA*), which codes for the major porin in *M. smegmatis.* This porin is important for the permeation of nutrients and antibiotics inside the cell. Previous studies showed that the deletion of this gene in *M. smegmatis* resulted in a reduced permeability to drugs such as β-lactams, fluoroquinolones and chloramphenicol [17-20]. The *mspA* gene is present in fast-growing mycobacteria only, and is not found in *M. tuberculosis* [21, 22].

**Table 2.**
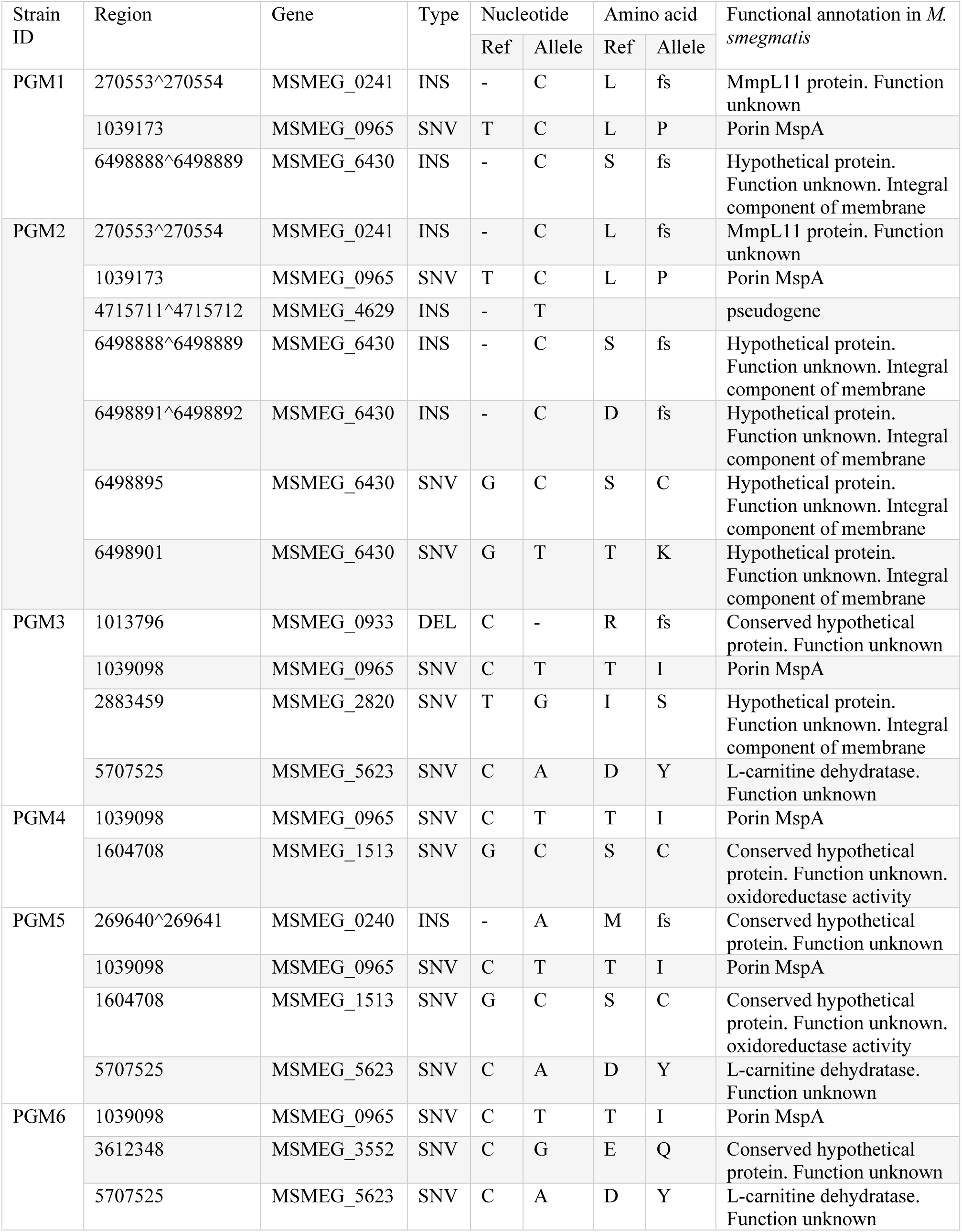
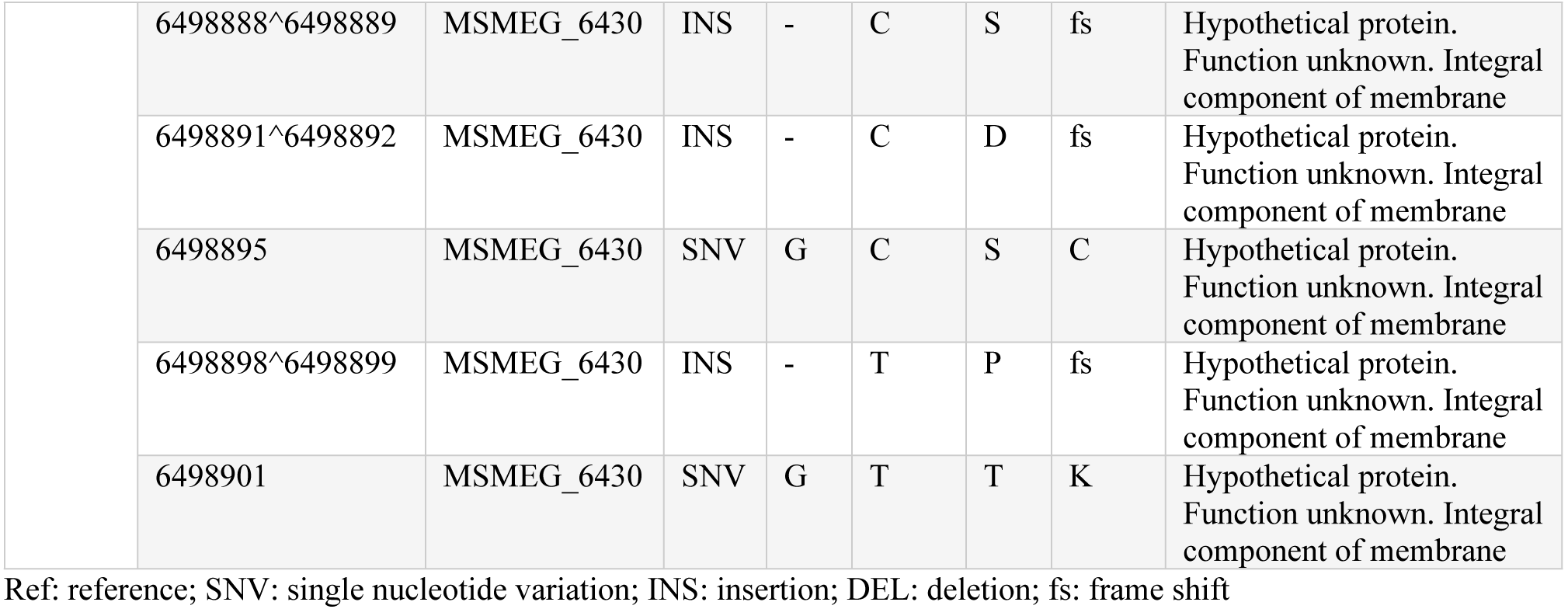
Mutations identified in each FP-11g resistant mutant

| Strain ID | Region | Gene | Type | Nucleotide |  | Amino acid |  | Functional annotation in *M. smegmatis* |
| --- | --- | --- | --- | --- | --- | --- | --- | --- |
|  |  |  |  | Ref | Allele | Ref | Allele |  |
| PGM1 | 270553^270554 | MSMEG_0241 | INS | - | C | L | fs | MmpL11 protein. Function unknown |
|  | 1039173 | MSMEG_0965 | SNV | T | C | L | P | Porin MspA |
|  | 6498888^6498889 | MSMEG_6430 | INS | - | C | S | fs | Hypothetical protein. Function unknown. Integral component of membrane |
| PGM2 | 270553^270554 | MSMEG_0241 | INS | - | C | L | fs | MmpL11 protein. Function unknown |
|  | 1039173 | MSMEG_0965 | SNV | T | C | L | P | Porin MspA |
|  | 4715711^4715712 | MSMEG_4629 | INS | - | T |  |  | pseudogene |
|  | 6498888^6498889 | MSMEG_6430 | INS | - | C | S | fs | Hypothetical protein. Function unknown. Integral component of membrane |
|  | 6498891^6498892 | MSMEG_6430 | INS | - | C | D | fs | Hypothetical protein. Function unknown. Integral component of membrane |
|  | 6498895 | MSMEG_6430 | SNV | G | C | S | C | Hypothetical protein. Function unknown. Integral component of membrane |
|  | 6498901 | MSMEG_6430 | SNV | G | T | T | K | Hypothetical protein. Function unknown. Integral component of membrane |
| PGM3 | 1013796 | MSMEG_0933 | DEL | C | - | R | fs | Conserved hypothetical protein. Function unknown |
|  | 1039098 | MSMEG_0965 | SNV | C | T | T | I | Porin MspA |
|  | 2883459 | MSMEG_2820 | SNV | T | G | I | S | Hypothetical protein. Function unknown. Integral component of membrane |
|  | 5707525 | MSMEG_5623 | SNV | C | A | D | Y | L-carnitine dehydratase. Function unknown |
| PGM4 | 1039098 | MSMEG_0965 | SNV | C | T | T | I | Porin MspA |
|  | 1604708 | MSMEG_1513 | SNV | G | C | S | C | Conserved hypothetical protein. Function unknown. oxidoreductase activity |
| PGM5 | 269640^269641 | MSMEG_0240 | INS | - | A | M | fs | Conserved hypothetical protein. Function unknown |
|  | 1039098 | MSMEG_0965 | SNV | C | T | T | I | Porin MspA |
|  | 1604708 | MSMEG_1513 | SNV | G | C | S | C | Conserved hypothetical protein. Function unknown. oxidoreductase activity |
|  | 5707525 | MSMEG_5623 | SNV | C | A | D | Y | L-carnitine dehydratase. Function unknown |
| PGM6 | 1039098 | MSMEG_0965 | SNV | C | T | T | I | Porin MspA |
|  | 3612348 | MSMEG_3552 | SNV | C | G | E | Q | Conserved hypothetical protein. Function unknown |
|  | 5707525 | MSMEG_5623 | SNV | C | A | D | Y | L-carnitine dehydratase. Function unknown |

|  |  |  |  |  |  |  |  |
| --- | --- | --- | --- | --- | --- | --- | --- |
| 6498888^6498889 | MSMEG_6430 | INS | - | C | S | fs | Hypothetical protein.<br>Function unknown. Integral<br>component of membrane |
| 6498891^6498892 | MSMEG_6430 | INS | - | C | D | fs | Hypothetical protein.<br>Function unknown. Integral<br>component of membrane |
| 6498895 | MSMEG_6430 | SNV | G | C | S | C | Hypothetical protein.<br>Function unknown. Integral<br>component of membrane |
| 6498898^6498899 | MSMEG_6430 | INS | - | T | P | fs | Hypothetical protein.<br>Function unknown. Integral<br>component of membrane |
| 6498901 | MSMEG_6430 | SNV | G | T | T | K | Hypothetical protein.<br>Function unknown. Integral<br>component of membrane |

Ref: reference; SNV: single nucleotide variation; INS: insertion; DEL: deletion; fs: frame shift

The second most frequent mutation (in 3 out of 6 mutants) detected corresponds to *MSMEG*_*5623* and *MSMEG*_*6430. MSMEG*_*5623* gene codes for an L-carnitine dehydratase homolog of unknown function in *M. smegmatis*, with no homolog in *M. tuberculosis* according to Tuberculist (http://svitsrv8.epfl.ch/tuberculist/). However, using the blast tool it was determined that this L-carnitine dehydratase homolog from *M. smegmatis* has 32% identity when aligned with the L-carnitine dehydratase from *M. tuberculosis* (*Rv3272)*, including Asp33 (conserved amino acid) that corresponds to the amino acid mutated in FP-11g resistant strains. *MSMEG*_*6430*, with a Thr to Lys substitution mutation detected in two mutants, is classified as a membrane protein that may have diverse functions in the cells, such as transportation of molecules through the membrane or serving as receptors for chemical signals [23, 24]. An Ile to Ser substitution mutation was also detected in *MSMEG*_*2820* which encodes for an unknown integral membrane protein. The mutations in diverse membrane proteins detected here may affect the transport of FP-11g across the membrane.

The remainder of detected mutations occurred at lower frequencies. *MSMEG*_*1513* (Ser to Cys substitution) codes for a hypothetical protein with no homolog in *M. tuberculosis* and has been classified as an oxidoreductase. *MSMEG*_*0241* (homolog of *Rv0202c*) with frame shift mutation detected encodes for MmpL11 (mycobacteria membrane protein large), a protein that belongs to a family of transporters and contribute to the cell wall biosynthesis in mycobacteria [25]. MmpL proteins family has been associated with drug resistance in *M. abscessus* and *M. tuberculosis* [26]. Frame shift mutation was also detected in *MSMEG*_*0240* (homolog of *Rv0201c*), which does not have a known function.

### Comparison of inhibition of MtbTopI versus DNA gyrase

The IC_50_ of FP-11g for inhibition of *E. coli* topoisomerase I relaxation activity has been reported as 0.48 μM [11]. In comparison, the IC_50_s for inhibition of human topoisomerase I relaxation activity and topoisomerase IIα decatenation activity were found to be both around 3.9 μM in the previous report [11]. Here we determined the IC_50_ for inhibition of MtbTopI and *Mtb* DNA gyrase by FP-11g (Fig 2). The IC_50_ for MtbTopI (0.24 μM) is ~100-fold lower than the IC_50_ for inhibition of *Mtb* DNA gyrase (31 μM).

**Fig 2.**
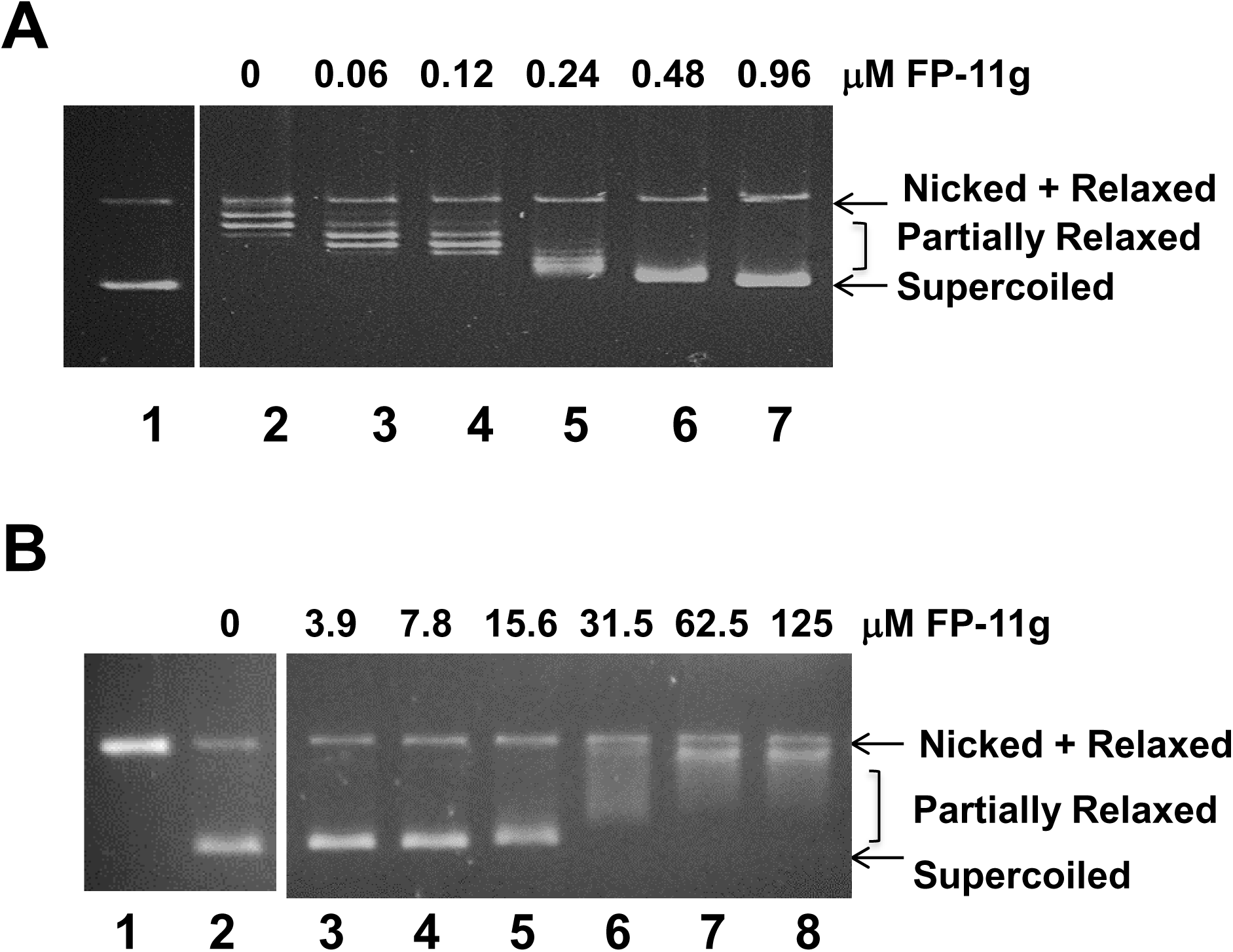
Comparison of inhibition of MtbTopI and DNA gyrase activities by FP-11g. **(A)** Inhibition of MtbTopI relaxation of negatively supercoiled DNA by increasing concentrations of FP-11g. **(B)** Inhibition of *Mtb* DNA gyrase supercoiling of relaxed DNA requires higher concentrations of FP-11g.

### FP-11g can interact directly with MtbTopI

To detect direct enzyme-inhibitor interaction, MtbTopI was immobilized on the CM5 chip surface and allowed to interact with FP-11g in the analyte solution. The SPR sensorgram (Fig 3) qualitatively showed that MtbTopI interacts directly with FP-11g.

**Fig 3.**
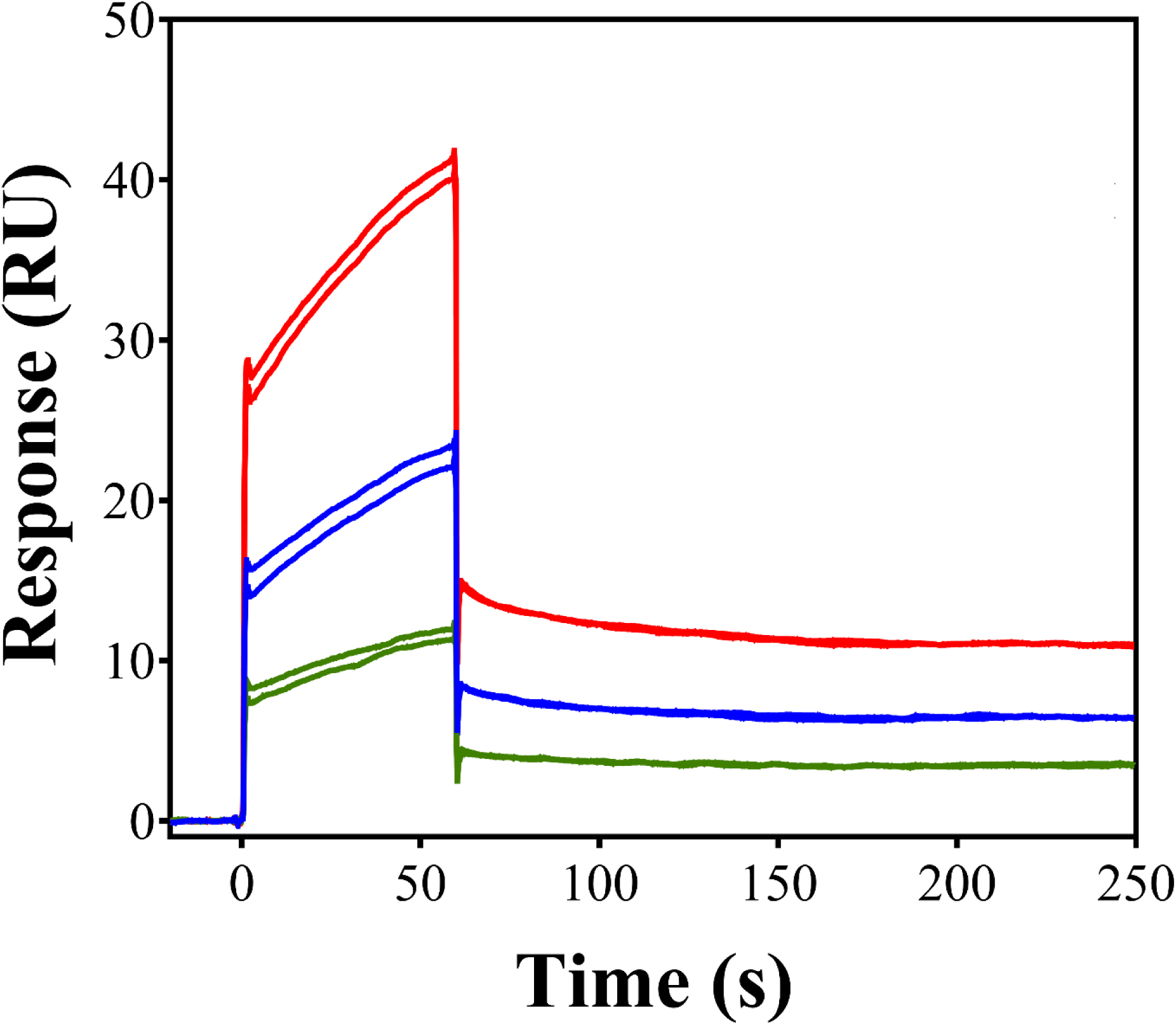
Qualitative evaluation of the binding of FP-11g with MtbTopI using SPR. SPR sensorgrams showing the direct binding of FP-11g with the immobilized MtbTopI. FP-11g was injected at 2.5 μM (green), 5 μM (blue), and 10 μM (red) in duplicate.

### FP-11g binds to DNA with intercalation

Since FP-11g has a planar tetracyclic ring scaffold (Scheme 1), it may bind to DNA by intercalation. Fig 4 shows the visible absorption spectra (Fig 4A) and fluorescence emission (Fig 4B) spectra of free and DNA-bound FP-11g. Binding of the compound to DNA results in a red shift of the maximum absorbance from 386 nm to 406 nm. Interestingly, upon binding to DNA, FP-11g has a pronounced induced visible band around 330 nm with a maximum absorbance at 334 nm. Binding of FP-11g to DNA also causes a red shift to the fluorescence emission spectrum of FP-11g (Fig 4B). The fluorescence intensity is slightly enhanced upon binding to DNA. Table 3 summarizes the optical properties of free and DNA-bound FP-11g.

**Fig 4.**
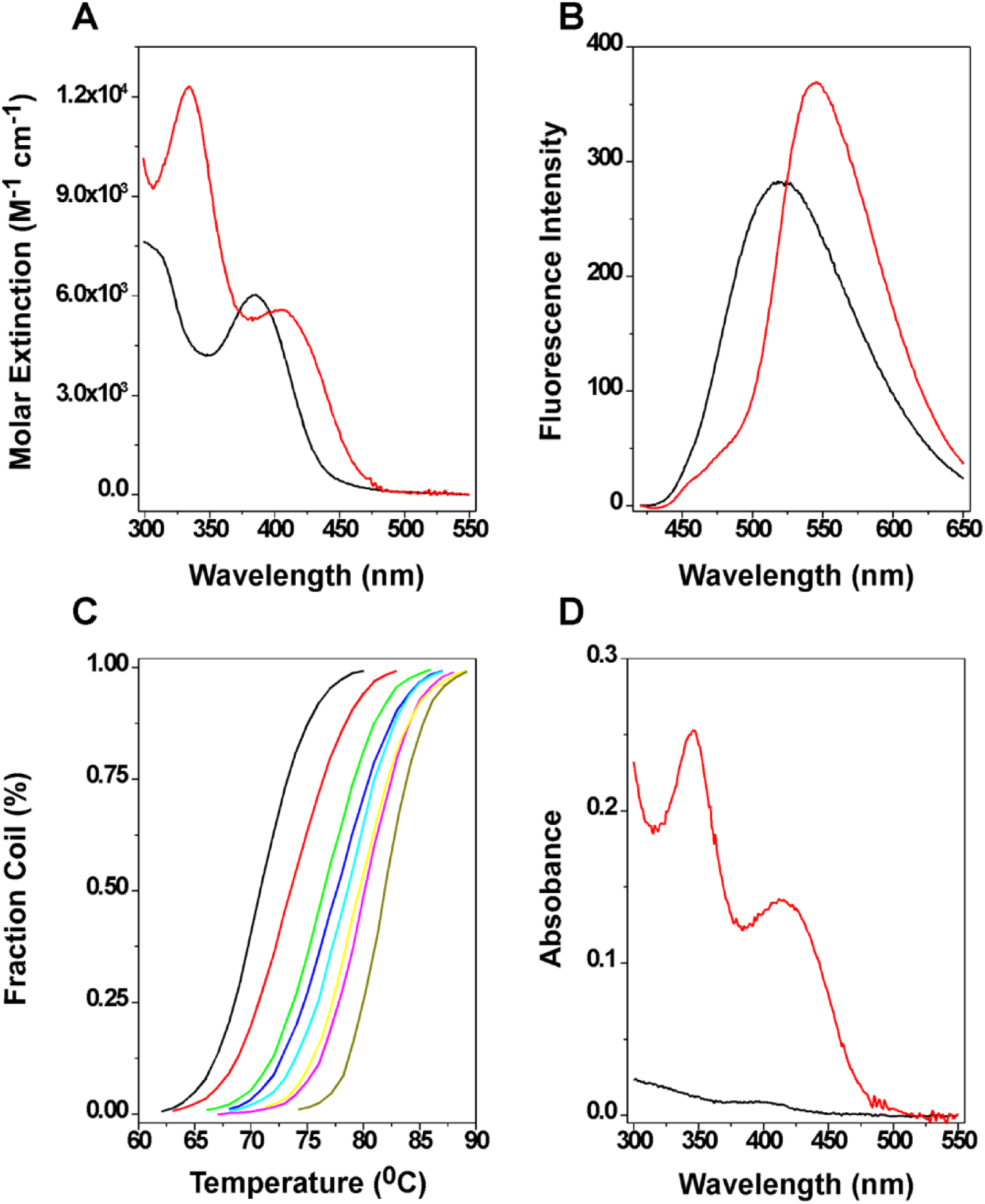
Binding of FP-11g to salmon testes DNA in 1xBPE buffer (6 mM Na_2_HPO_4_, 2 mM NaH_2_PO_4_, and 1 mM EDTA, pH 7). **(A)** Visible absorption spectra of free (black line) and DNA bound (red line). **(B)** Fluorescence emission spectra of free (black line) and DNA bound (red line). The fluorescence emission spectra were recorded with λ_em_ = nm. **(C)** DNA UV melting of salmon testes DNA in the presence of various concentrations of FP-11g. The FP-11g concentrations are 0, 2, 5, 7.5, 10, 15, 20, and 40 μM from left to right. **(D)** The DNA dialysis assay was performed as described in the Methods. Visible absorption spectra of FP-11g inside the dialysis bag (red line) and outside dialysis bag (black line).

**Table 3.**
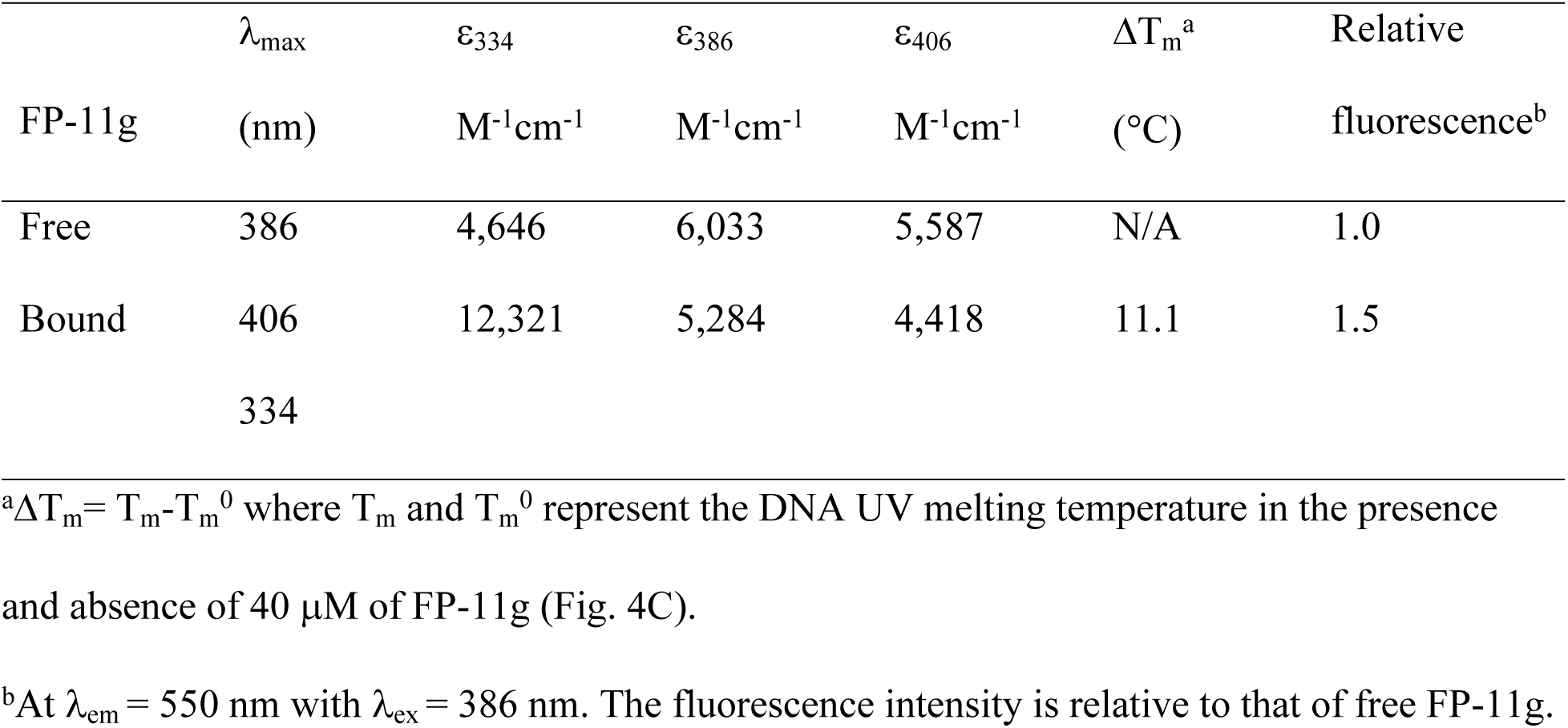
Optical properties of FP-11g in the presence and absence of ST DNA in 1×BPE buffer

| | $\lambda_{\text{max}}$ | $\epsilon_{334}$ | $\epsilon_{386}$ | $\epsilon_{406}$ | $\Delta T_m^a$ | Relative |
| --- | --- | --- | --- | --- | --- | --- |
| FP-11g | (nm) | $\text{M}^{-1}\text{cm}^{-1}$ | $\text{M}^{-1}\text{cm}^{-1}$ | $\text{M}^{-1}\text{cm}^{-1}$ | (°C) | fluorescence<sup>b</sup> |
| Free | 386 | 4,646 | 6,033 | 5,587 | N/A | 1.0 |
| Bound | 406 | 12,321 | 5,284 | 4,418 | 11.1 | 1.5 |
|  | 334 |  |  |  |  |  |

<sup>a</sup> $\Delta T_m = T_m - T_m^0$ where $T_m$ and $T_m^0$ represent the DNA UV melting temperature in the presence and absence of 40 $\mu\text{M}$ of FP-11g (Fig. 4C).
<sup>b</sup>At $\lambda_{\text{em}} = 550$ nm with $\lambda_{\text{ex}} = 386$ nm. The fluorescence intensity is relative to that of free FP-11g.

Fig 4C shows the DNA UV melting profiles in which different concentrations of FP-11g were added to the same concentration of ST DNA. In the absence of FP-11g, the UV melting temperature of ST DNA (Tm) was determined to be 70.8°C. In the presence of saturated FP-11g (40 μM), the DNA UV melting temperature was increased to 81.9°C, indicating that FP-11g tightly binds to ST DNA. In this study, we also carried out a dialysis binding assay in which 75 μM (bp) ST DNA was extensively dialyzed against a large excess of 1 μM of FP-11g. Fig 4D shows our results. After 72 hours of dialysis, the concentration of FP-11g outside the dialysis bag was not significantly changed. In contrast, the concentration of FP-11g inside the dialysis bag where the ST DNA was located was increased to 25 μM. The DNA binding constant of FP-11g was estimated to be 1×10^6^ M^−1^ in 1XBPE.

Fig 5 shows the results of ligation of nicked circular DNA in the presence of different concentrations of FP-11g by T4 DNA ligase. Change in DNA linking number was observed at 2 μM. Concentrations of FP-11g at 5 and 10 μM drove the plasmid DNA template into (-) supercoiled DNA. This result suggests that FP-11g can act as a DNA intercalator at these higher concentrations. The DNA intercalation of FP-11g at the observed IC_50_s could be the basis for the inhibition of DNA gyrase and human topoisomerases [11], but could not be the sole basis of inhibition of bacterial topoisomerase I observed at 10-fold lower IC_50_s.

**Fig 5.**
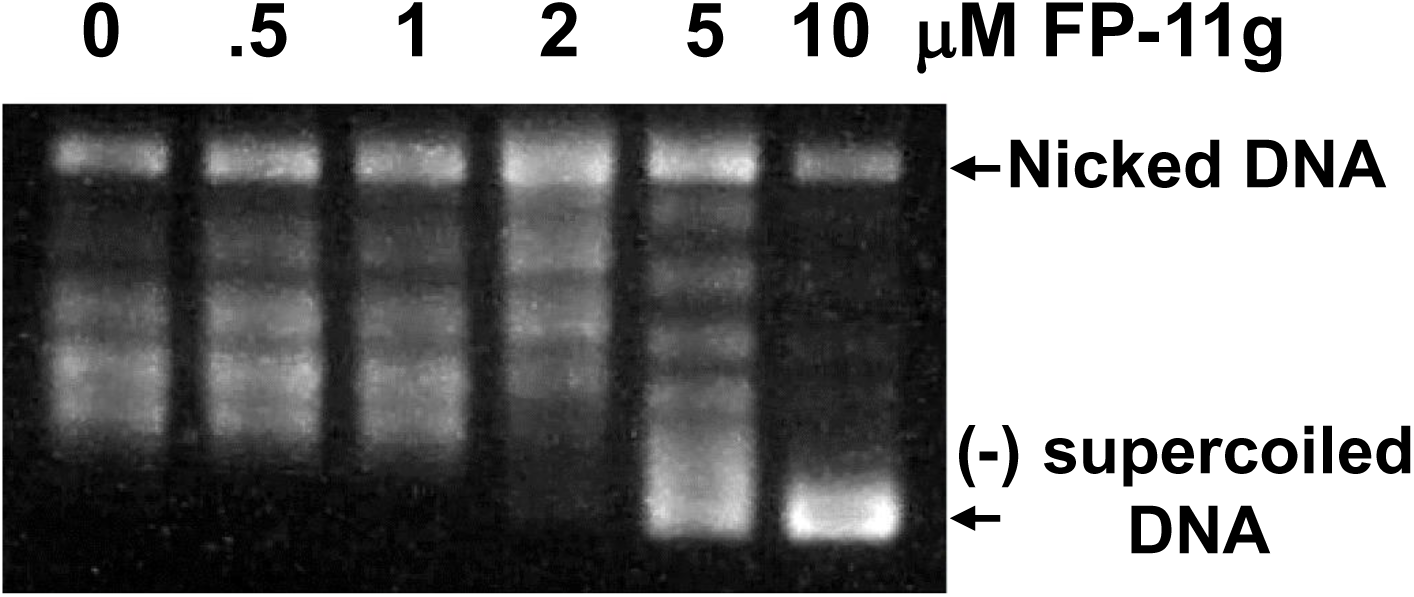
Products of DNA ligation in the presence of FP-11g. DNA ligation of nicked pAB1 plasmid DNA by T4 DNA ligase in the presence of indicated concentrations of FP-11g were performed as described under Methods.

## Discussion

In previous studies, the FP-11g compound has been proposed as an antimycobacterial agent active against *M. tuberculosis* (MIC = 2.5 μM) [11]. Here we demonstrate its activity against the pathogenic NTM *M. abscessus.* Compounds active against both species would be useful in clinical settings since in some cases the symptoms and clinical manifestations of infections caused by these two organisms are difficult to differentiate, consequently empiric treatment effective for both organisms is required. Additionally, patients suffering from cystic fibrosis or other immunosuppressive condition can be infected by different mycobacteria at the same time. In fact, co-infections with *M. abscessus* and *M. tuberculosis* have been reported [27, 28]. The strong bactericidal effect of FP-11g against *M. smegmatis* supports investigation of this compound and its derivatives for its potential use as a broad antimycobacterial agent. The finding of the growth inhibitory effect of FP-11g on *M. abscessus* is encouraging because of the lack of response of NTM clinical strains and subspecies of *M. abscessus* to nearly all of the current antibiotics.

No specific mutations found in FP-11g *M. smegmatis* resistant mutants could be definitively associated with the proposed mechanism of action of this compound, but all of them may play a role in drug resistance. Expression of *M. smegmatis* MspA porin in *M. tuberculosis* promotes not only cell growth but also antibiotic susceptibility [29]. To date, porin genes have not yet been identified in *M. tuberculosis*, even though some studies suggest the presence of these structures in these organisms [22]. The mutation in MspA porin is likely to be the first mechanism of resistance developed by *M. smegmatis*, and hence the common mutation was found. Changes in the porin may interrupt the drug transportation into the cell and support cell survival. Since this porin is present in fast growing mycobacteria, it would be of significance in the treatment of atypical pathogenic mycobacteria including *M. abscessus* studied here [30].

Characterization of integral membrane proteins has been always a challenge due to the difficulty for the expression, solubility and crystallization of these proteins. Most of the information has been obtained through computational approaches, which predict the structural conformation of a protein based on the primary sequence and classify the different types of transmembrane proteins [31, 32]. Transmembrane proteins could be associated with mechanisms of resistance to drugs that are not targeting them directly. Integral component of membrane are channels through the cellular membrane that transport amino acids, lipids, coenzymes, carbohydrates, nucleotides and other metabolites [33]. Mutations in integral membrane proteins detected here may affect drug transportation through the membrane. Variation in channels that transport molecules through the cell wall will affect the intake of molecules including drugs and metabolites. These integral components of membrane could also be non-characterized efflux pumps that may be extruding the drug from the organism and support the drug resistance.

In this study we detected the MmpL11 mutation in two resistant mutants. Previous studies with *M. tuberculosis* strains harboring mutations in MmpL proteins showed that all of the mutant strains retained general drug susceptibility to diverse antibacterial agents. This suggests that MmpL proteins do not play a direct role in drug resistance [34]. Nonetheless, recent studies have shown that the extended RND permease superfamily in *M. tuberculosis* include both RND multidrug efflux transporters and members of the MmpL family [35]. In *M. abscessus*, MmpL has been associated with drug resistance through efflux pumps [36]. Additionally, mutations on MmpL11 proteins has been associated with impairment of biofilm formation in *M. smegmatis.* The absence of this gene has generated a reduced permeability to antimicrobial agents [37, 38], which is evidence that this mutation may play an important role in FP-11g resistance.

Remarkably, we did not detect mutations in the *topA* gene in the resistant mutant isolates. This might be due to FP-11g acting as an unconventional catalytic inhibitor. These types of inhibitors are described as compounds that bind to the DNA as well as to the topoisomerase enzyme to inhibit the topoisomerase activity [39]. Mutations that affect the essential activity of topoisomerase I could compromise cell growth to a significant extent and would less likely be selected for resistance than the mutations detected here that can limit compound transport into the cell. It is also possible that due to its interaction with DNA, FP-11g inhibits multiple DNA binding proteins in its mode of action, including DNA gyrase as observed here. Further studies are required to identify other analogs of FP-11g that can be more selective in targeting topoisomerase I activity in its antimycobacterial mechanism of action. This would require enhancing the interaction of the inhibitor with the MtbTopI enzyme, and reducing the dependence of inhibition on DNA binding.

## Acknowledgments

We thank Dr. DeEtta Mills and Christina Burns from the International Forensic Research Institute at FIU for support in WGS.

## Funding

This research was supported by NIH grant R01 AI069313 (to YT). Experimental SPR sensorgrams were measured using Biacore T200 SPR instrument available in Biacore Molecular Interaction Shared Resource (BMISR) facility at Georgetown University. The BMISR is supported by NIH grant P30CA51008. REU participant RB was supported by the NSF-REU Site Grant CHE1560375 to FIU. The funders had no role in study design, data collection and analysis, decision to publish, or preparation of the manuscript.

---

**Table S1.** Summary of mutations associated with FP-11g resistance in *M. smegmatis*

| Gene | Homolog in *Mtb* | Region | Freq | Type | Nucleotide |  | Amino acid |  | Functional annotation in *M. smegmatis* | Functional annotation in *M. tuberculosis* |
| --- | --- | --- | --- | --- | --- | --- | --- | --- | --- | --- |
|  |  |  |  |  | Ref | Allele | Ref | Allele |  |  |
| MSMEG_0965 | No | 1039098 | 4 | SNV | C | T | T | I | Porin MspA | N/A |
| MSMEG_0965 | No | 1039173 | 2 | SNV | T | C | L | P | Porin MspA | N/A |
| MSMEG_2820 | No | 2883459 | 1 | SNV | T | G | I | S | Hypothetical protein. Function unknown. Integral component of membrane | N/A |
| MSMEG_6430 | No | 6498888^6498889 | 3 | INS | - | C | S | fs | Hypothetical protein. Function unknown. Integral component of membrane | N/A |
| MSMEG_6430 | No | 6498891^6498892 | 2 | INS | - | C | D | fs | Hypothetical protein. Function unknown. Integral component of membrane | N/A |
| MSMEG_6430 | No | 6498895 | 2 | SNV | G | C | S | C | Hypothetical protein. Function unknown. Integral component of membrane | N/A |
| MSMEG_6430 | No | 6498898^6498899 | 1 | INS | - | T | P | fs | Hypothetical protein. Function unknown. Integral component of membrane | N/A |
| MSMEG_6430 | No | 6498901 | 2 | SNV | G | T | T | K | Hypothetical protein. Function unknown. Integral component of membrane | N/A |
| MSMEG_0241 | Rv0202c | 270553^270554 | 2 | INS | - | C | L | fs | MmpL11 protein. Function unknown | unknown/cell wall and cellular processes |
| MSMEG_5623 | No | 5707525 | 3 | SNV | C | A | D | Y | L-carnitine dehydratase. Function unknown | N/A |
| MSMEG_0933 | Rv0486 | 1013796 | 1 | DEL | C | - | R | fs | Conserved hypothetical protein. Function unknown | *mshA* gene: Glycosyltransferase |
| MSMEG_0240 | Rv0201c | 269640^269641 | 1 | INS | - | A | M | fs | Conserved hypothetical protein. Function unknown | conserved hypothetical /unknown |
| MSMEG_1513 | No | 1604708 | 2 | SNV | G | C | S | C | Conserved hypothetical protein. Function unknown. oxidoreductase activity | N/A |
| MSMEG_4629 |  | 4715711^4715712 | 1 | INS | - | T |  |  | pseudogene | N/A |
| MSMEG_3552 | No | 3612348 | 1 | SNV | C | G | E | Q | Conserved hypothetical protein. Function unknown | N/A |

Freq: frequency; SNV: single nucleotide variation; INS: insertion; DEL: Deletion; Ref: reference; fs: frame shift

**Scheme 1.**
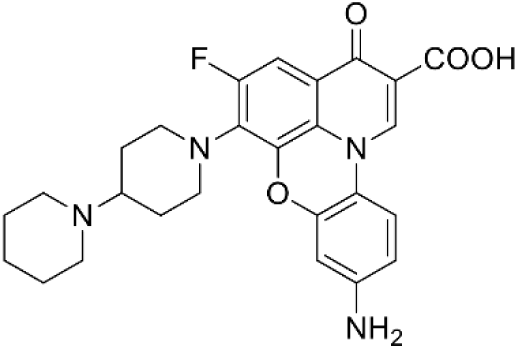
Structure of FP-11g

